# Sex-Specific Function and Morphology of the Anterior Cruciate Ligament During Skeletal Growth in a Porcine Model

**DOI:** 10.1101/2021.05.10.442986

**Authors:** Danielle Howe, Stephanie G. Cone, Jorge A. Piedrahita, Bruce Collins, Lynn A. Fordham, Emily H. Griffith, Jeffrey T. Spang, Matthew B. Fisher

**Affiliations:** Joint Department of Biomedical Engineering, North Carolina State University and the University of North Carolina- Chapel Hill; Raleigh, NC 27695; Comparative Medicine Institute, North Carolina State University; Raleigh, NC 27695; Department of Biomedical Engineering, University of Wisconsin- Madison; Madison, WI 53706; Department of Molecular Biomedical Sciences, North Carolina State University; Raleigh, NC 27695; Department of Animal Science, North Carolina State University; Raleigh, NC 27695; Department of Radiology, University of North Carolina- Chapel Hill; Chapel Hill, NC 27599; Department of Statistics, North Carolina State University; Raleigh, NC 27695; Department of Orthopaedics, University of North Carolina- Chapel Hill; Chapel Hill, NC 27599

**Keywords:** anterior cruciate ligament, knee, skeletal growth, porcine model

## Abstract

Pediatric anterior cruciate ligament (ACL) injuries are on the rise, and females experience higher ACL injury risk than males during adolescence. Studies in skeletally immature patients indicate differences in ACL size and joint laxity between males and females after the onset of adolescence. However, functional data regarding the ACL and its anteromedial and posterolateral bundles in the pediatric population remain rare. Therefore, this study uses a porcine model to investigate the sex-specific morphology and function of the ACL and its bundles throughout skeletal growth. Hind limbs from male and female Yorkshire pigs aged early youth to late adolescence were imaged using magnetic resonance imaging to measure the size and orientation of the ACL and its bundles, then biomechanically tested under anterior-posterior drawer using a robotic testing system. Joint laxity decreased (p<0.001) while joint stiffness increased (p<0.001) throughout skeletal growth in both sexes. The ACL was the primary stabilizer against anterior tibial loading in all specimens, while the functional role of the anteromedial bundle increased with age (p<0.001), with an earlier shift in males. ACL and posterolateral bundle cross-sectional area and ACL and anteromedial bundle length were larger in males than females during adolescence (p<0.01 for all), while ACL and bundle sagittal angle remained similar between sexes. Additionally, *in situ* ACL stiffness correlated with cross-sectional area across skeletal growth (r^2^=0.75, p<0.001 in males and r^2^=0.64, p<0.001 in females), but not within age groups. This study has implications for age and sex-specific surgical intervention strategies and suggests the need for human studies.

## Introduction

Anterior cruciate ligament (ACL) injuries and surgical reconstruction are increasingly common in the skeletally immature population,^1; 2^ and approximately one third of pediatric patients experience a second ACL injury after surgical reconstruction.^3^ There appears to be an interaction between age and sex regarding ACL injury risk, such that females experience a greater risk than males during adolescence, but not during childhood or adulthood.^4^ These data point to the challenge of restoring ACL function in pediatric patients, as the demand placed on the normal ACL or replacement graft may vary with age and sex.

Studies have examined sex-specific morphology and function of the ACL throughout skeletal growth in humans to try to explain age- and sex-specific injury rates. The ACL is similar in size between males and females during childhood, but becomes larger in length and cross-sectional area (CSA) in males after the onset of adolescence.^5–8^ Additionally, anterior-posterior (AP) knee laxity decreases throughout skeletal growth with similar values between males and females during childhood.^9–11^ During adolescence^9–11^ and adulthood,^12^ AP knee laxity appears greater in females than males. Although greater passive AP knee laxity has been linked to ACL injury risk,^12–14^ it is unclear that differences of only ~1 mm observed in these studies^9–12^ can account for the difference in injury rate between sexes. However, age and sex-based differences in joint loading and kinematics also appear during dynamic activities.^15; 16^ Additionally, the biomechanical properties of the pediatric ACL have been measured directly in one recent study from a small set (n=5) of pediatric cadaveric specimens.^17^ Under uniaxial tension, the pediatric ACL exhibited lower mechanical properties^17^ compared to those in the adult ACL.^18^ However, the small sample size of this subset limits further age and sex-specific comparisons.

Despite the usefulness of the aforementioned studies, there remains an incomplete understanding of the complex function of the ACL in the skeletally immature population. The ACL can be divided into two functional bundles: the anteromedial (AM) and posterolateral (PL) bundles. These two bundles vary in their insertion to the lateral femoral condyle and tibial plateau and have a complementary function in adults. For example, the relative contribution of the AM bundle increases with increasing knee flexion in humans, while the PL bundle plays a larger role near full knee extension.^19–21^ However, the functional maturation of the AM and PL bundles in skeletally immature humans has not been studied, primarily due to a dearth of pediatric cadaveric specimens. Consequently, there is a need for preclinical models of the pediatric ACL.

Our group and others have used the porcine model to study the skeletally immature ACL^22–27^ and sex-specific ACL behavior.^28; 29^ The porcine model is widely used for musculoskeletal applications^30^ and is a good model to study function of the mature ACL.^31; 32^ The female porcine ACL increases in length, CSA, and angular orientation throughout skeletal growth^22; 23^, similar to humans.^5–8; 33^ These studies also revealed age-specific differences in growth and function of the AM and PL bundles.^23^ However, these findings leave open the question of whether functional maturation of the ACL and its bundles is sex-specific.

Therefore, the objective of this study was to investigate the morphology and function of the ACL and its bundles as a function of sex throughout skeletal growth in the porcine model. To accomplish this, we imaged stifle (knee) joints from male and female pigs at five age groups throughout skeletal growth using magnetic resonance imaging (MRI) to assess ACL and bundle morphology. Then, in the same joints, we used a robotic testing system to measure joint kinematics and ACL and bundle force contributions under applied AP loads.

## Methods

### Specimen Collection

A total of 60 hind limbs were collected from male and female Yorkshire cross-breed pigs at 1.5 (early juvenile), 3 (juvenile), 4.5 (early adolescent), 6 (adolescent), and 18 (late adolescent) months of age (n=6 animals/age/sex, 1 limb per animal) (Fig. 1A). All animals were bred at the North Carolina State University Swine Educational Unit, and all experimental protocols were approved by the North Carolina State University Institutional Animal Use and Care Committee. Immediately following euthanasia, hind limbs were disarticulated, and stifle (knee) joints were isolated. Joints were wrapped in saline-soaked gauze and stored at −20°C. Data from female specimens has been previously reported.^22–24^

**Figure 1.**
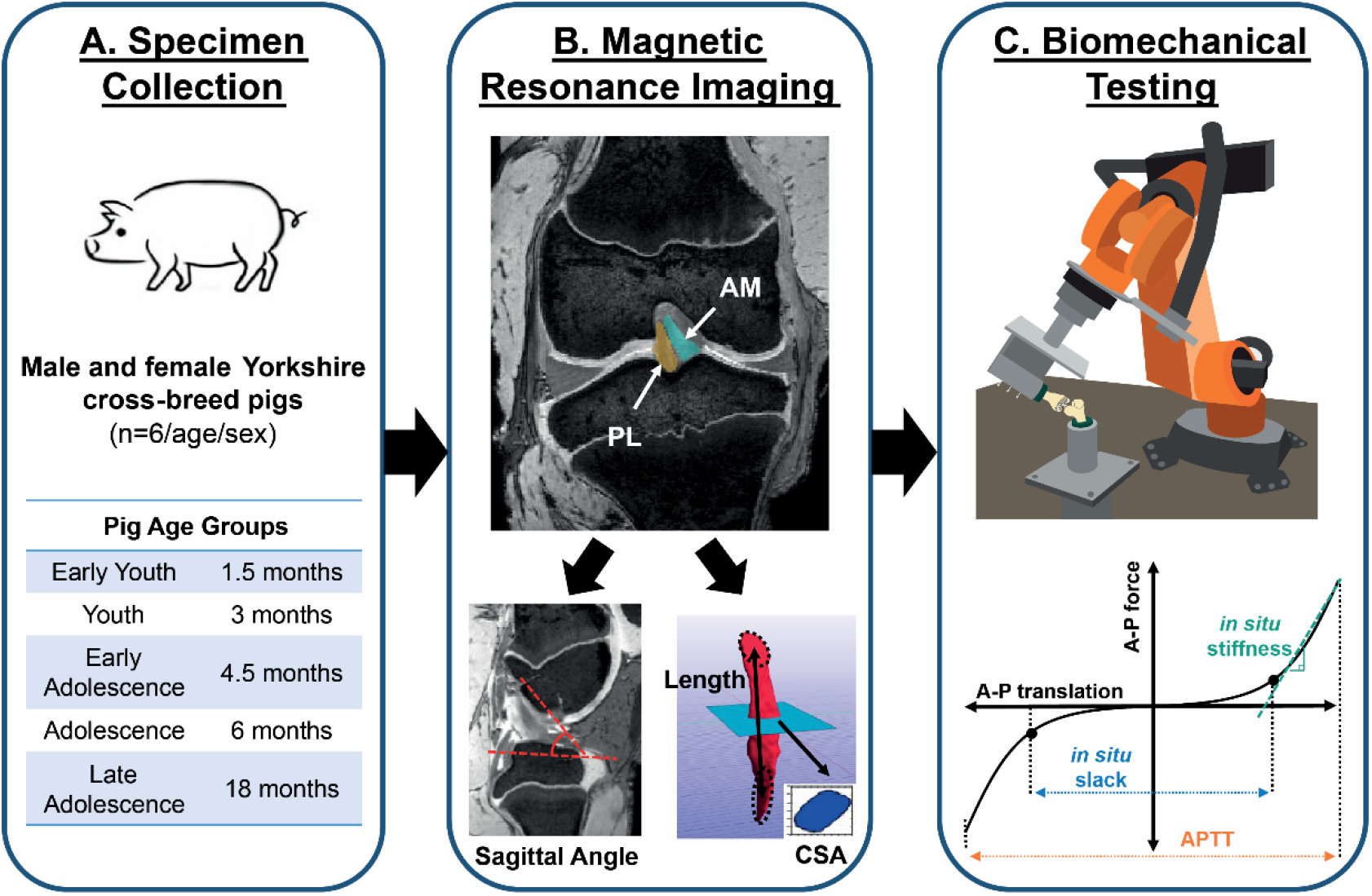
Experimental methods overview. (A) Hind limbs were collected from male and female pigs at the age groups listed (n=6/age/sex). (B) Magnetic resonance images were used to measure the length, cross-sectional area (CSA), and angular orientation of the ACL and its anteromedial (AM) and posterolateral (PL) bundles. (C) Biomechanical testing was performed with a robotic testing system equipped with a universal force-moment sensor. Anterior-posterior (A-P) loads were applied to each joint, and A-P force translation curves were used to determine the total A-P tibial translation (APTT), *in situ* joint slack, and *in situ* joint stiffness. ACL and bundle force contributions were measured at peak anterior translation.

### Magnetic Resonance Imaging

Stifle joints were thawed at room temperature and imaged at full extension (30°-40° of flexion). Images were acquired with a 7.0-T Siemens Magnetom MRI scanner (Siemens Healthineers, Erlangen, Germany) using a 28-channel knee coil, and double-echo steady state sequence (DESS, flip angle: 25°, TR: 17ms, TE: 6s, voxel size: 0.42×0.42×0.4 mm, no gap between slices). Joints were stored at −20°C after imaging.

Images were analyzed using commercial software (Simpleware ScanIP, Synopsys, Mountain View, CA, USA) (Fig. 1B). The angular orientation of the ACL, AM bundle, and PL bundle was measured relative to the tibial plateau in the sagittal plane, as previously described.^22; 23^ The ACL and its AM and PL bundles were manually segmented from each image, then exported as surface models. Models were analyzed in Matlab (MathWorks, Natick, MA, USA), as previously described.^23^ Briefly, the CSA of each ACL, AM bundle, and PL bundle was calculated as the average CSA of the central 50% of each tissue, perpendicular to its primary axis. The length of each tissue was calculated as the distance between the centroid of its femoral and tibial insertion site.

### Biomechanical Testing

Biomechanical testing was performed using a robotic testing system (KR300 R2500, KRC4, Kuka, Shelby Charter Township, MI, USA) equipped with a universal force-moment sensor (Omega160 IP65, ATI Industrial Automation, Apex, NC, USA), as previously described.^23; 24^ Systems were integrated and controlled using the simVitro software knee module (Cleveland Clinic, Cleveland, OH, USA). Stifle joints were thawed at room temperature to be prepared for robotic testing. The femur and tibia were fixed in custom molds using a fiberglass reinforced epoxy (Everglass, Evercoat, Cincinnati, OH, USA).

For each joint, the femur was fixed rigidly to the testing platform, and the tibia was fixed to the robotic testing system using custom clamps (Fig. 1C). Joints were lightly wrapped in saline soaked gauze throughout testing. An anatomic coordinate system was established for each stifle using a 3D digitizer (G2X MicroScribe, Amherst, VA, USA). Each joint was flexed from 40° of flexion (approximately full extension) to 90° of flexion while minimizing forces in the remaining five degrees of freedom to determine a passive path. AP forces were then applied to the tibia at 60° of flexion with internal rotation and flexion angle fixed and minimizing forces in the remaining three degrees of freedom. Preliminary tests on female joints showed that fixing internal rotation did not impact joint kinematics under AP drawer (Table S1). Loads were scaled to the femoral cortical CSA for each age group (20 N, 40 N, 80 N, 100 N, 140 N for 1.5, 3, 4.5, 6, and 18 month-old joints, respectively). The kinematic paths under the applied loads were recorded and repeated while recording the forces. Next, the joint capsule was dissected to visualize the ACL. Preliminary data showed that the capsule alone contributed minimally to anterior drawer (Table S2). The AM and PL bundles were separated with a blunt instrument at the tibial insertions, and the AM bundle was transected by cutting anterior to the instrument. To engage the PL bundle, forces were applied to the AM bundle deficient joint and a new set of kinematics was recorded. The kinematic paths from the intact and AM bundle deficient states were repeated for the AM bundle deficient knee while recording forces. Next, the PL bundle was transected, and the kinematic paths were repeated while recording forces in the ACL deficient state.

AP force-translation curves were plotted for each joint, with data collected and plotted at increments of 20% of the maximum applied force (Fig. 1C, Fig. 2A). The maximum anterior-posterior tibial translation (APTT) was measured and normalized to the AP tibial width of each joint to assess joint laxity. To assess sub-maximal joint behavior, *in situ* joint slack and stiffness were calculated using Matlab, as previously described.^24; 34^ Biphasic curves consisting of an exponential region and a linear region were fitted to the anterior and posterior portions each force-translation curve. Anterior and posterior engagement points were determined as the points of maximum curvature.^34; 35^ *In situ* joint slack was defined as the distance between the anterior and posterior engagement points. *In situ* joint stiffness was defined as the slope of a linear curve fitted from the anterior engagement point to maximum anterior translation.

**Figure 2.**
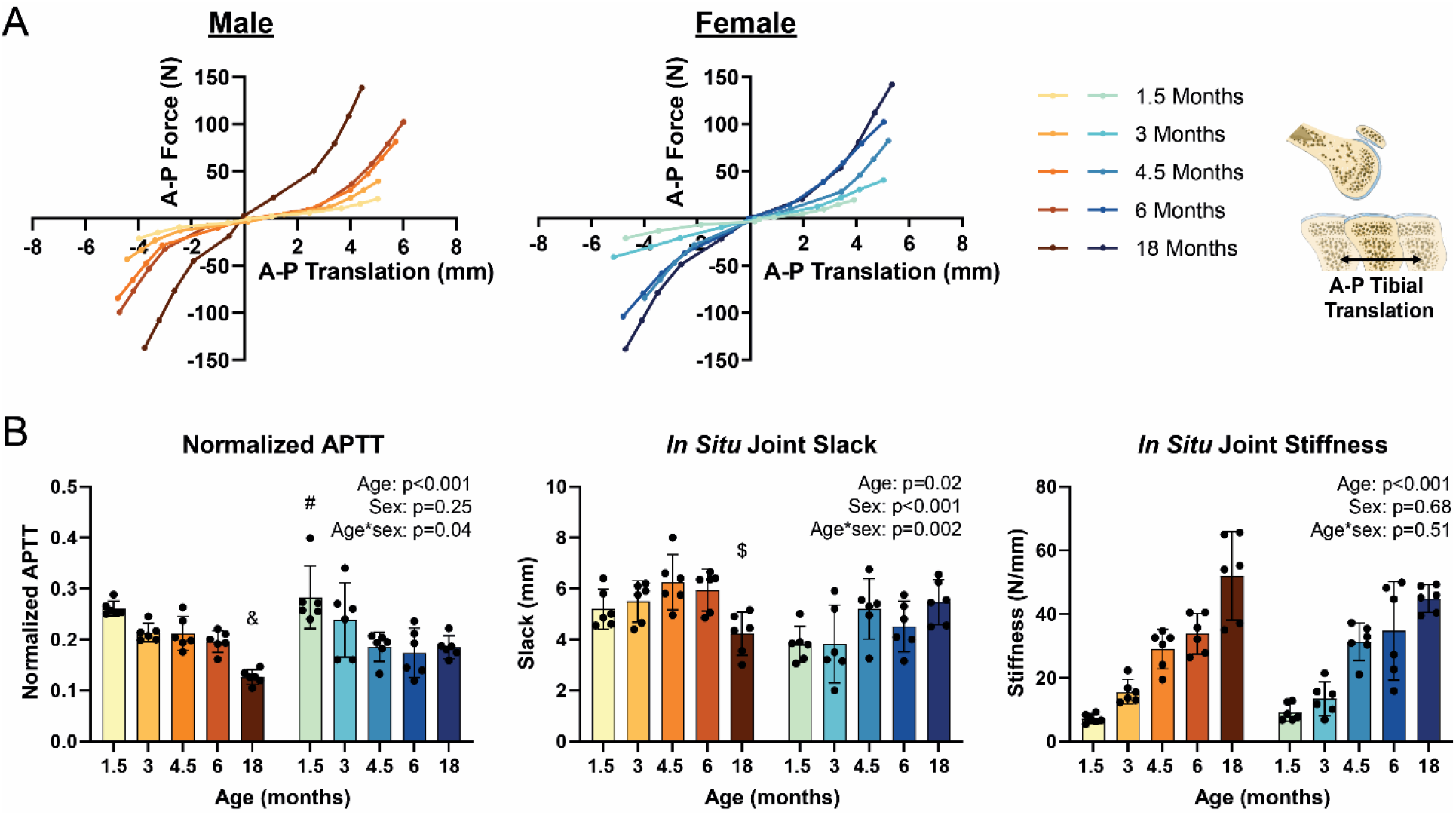
Knee joint function shifted throughout skeletal growth. (A) In response to an anterior-posterior (A-P) load, mean force-translation curves for male and female joints became steeper throughout growth, with a sizeable shift from 6 to 18 months in males but a more gradual shift between ages in females. (B) Quantitative parameters of joint function included normalized anterior-posterior tibial translation (APTT), joint *in situ* slack, and joint *in situ* stiffness. Throughout skeletal growth, normalized APTT decreased while joint stiffness increased in both males and females. Joint slack showed little change with age, apart from a significant decrease from 4.5 to 18 months in males. Data presented as mean ± 95% confidence interval with main effects from two-way ANOVA shown in graph corner. & indicates p<0.01 compared to 1.5, 3, 4.5 months. # indicates p<0.01 compared to 4.5, 6, and 18 months. $ indicates p<0.01 compared to 4.5 months.

The force contributions of the ACL, AM bundle, and PL bundle during anterior drawer were calculated using the principle of superposition.^36; 37^ The anterior force carried by the ACL was calculated as a percentage of the total anterior force in the knee joint under maximum anterior translation. The anterior forces carried by the AM and PL bundles were calculated as a percentage of the anterior force in the ACL under maximum anterior translation. *In situ* slack and stiffness of the ACL and AM bundle were calculated by plotting anterior tissue force versus anterior joint translation from the intact kinematics. PL bundle *in situ* slack and stiffness was calculated by plotting anterior tissue force versus anterior joint translation from the AM bundle deficient kinematics.

### Statistical Analysis

Statistical analysis was performed in Prism (GraphPad, San Diego, CA, USA). Outlier testing was performed on all data sets using the ROUT method (Q=10%). One 18 month male specimen was identified as an outlier in APTT, normalized APTT, and slack length and was replaced. Normality of all data sets was confirmed. Joint and tissue functional and structural parameters were assessed using two-way ANOVA and Tukey’s post hoc test (α=0.05), with age and sex as main effects. To compare bundle function, the difference in AM and PL bundle percent contribution was calculated for each specimen. This difference in bundle contribution was analyzed with two-way ANOVA with one sample t-tests comparing mean values to zero to test for significant, non-zero differences in bundle contribution. Linear regression analyses were performed for ACL *in situ* stiffness versus CSA for each sex first pooled across ages, then for each age group. Linear regression analyses were also performed for AM and PL bundle *in situ* stiffness versus CSA separated by sex and estimated pubertal status^38^—pre-pubertal (1.5-4.5 months) and post-pubertal (6 and 18 months). Figures include individual data points overlaid with bars representing mean ± 95% confidence interval.

## Results

### Joint Function

Qualitatively, the average AP force-translation curves of male and female stifle joints became progressively steeper throughout skeletal growth (Fig. 2A). There was a large shift of average force-translation curves between 6 and 18 months in males, compared to a more gradual shift between collected timepoints in females. Comparing between males and females at each age (Fig. S1), average curves appeared similar from 1.5-4.5 months, but at 6 months, the male force-translation curve extended out wider than females. However, at 18 months, force-translation curves appeared narrower and steeper for male joints.

Joint behavior was quantitatively evaluated through normalized APTT, joint *in situ* slack, and joint *in situ* stiffness (Fig. 2B). Mean normalized APTT decreased from 1.5-18 months by half in males and by one third in females (p<0.001, Table S3). In females, this decrease occurred primarily between 3-6 months while in males, the largest decrease occurred between 6-18 months. Normalized APTT did not differ significantly between the sexes across all ages (p=0.25). Normalized APTT was within 15% between age-matched males and females from 1.5-6 months, and males had 38% lower values than females at 18 months, though this difference was not statistically significant (p=0.19).

*In situ* joint slack length varied in a sex-specific manner (p=0.002 for age-sex interaction, Fig. 2B, Table S5). For males, average joint slack showed little change from 1.5-6 months, but decreased significantly by 18 months (38%, p=0.018 between 4.5 and 18 months). Female joint slack did not significantly differ with age. Across all ages, joint slack was 17% greater in males than females (p<0.001); although, there were not significant differences between the sexes within age groups.

*In situ* joint stiffness increased from 1.5-18 months in males and females by approximately 7-fold and 5-fold, respectively (p<0.001, Fig 2B, Table S6). In males, the largest increases in joint stiffness were observed from 3-4.5 months (60%) and 6-18 months (42%). In females, the largest increase in joint stiffness was observed from 3-4.5 months (80%), with only a 25% increase from 6-18 months. Overall, sex did not significantly impact joint stiffness (p=0.68).

### ACL and Bundle Function

Across all joints, the ACL carried 75-112% of the applied anterior force, with no significant impact of age or sex (Fig. 3A, Table S7). However, within the ACL, the anterior force distribution between the AM and PL bundles showed age and sex dependent shifts (Fig. 3B, C, Table S8). Both bundles carried substantial force during youth; mean AM and PL bundle contributions to the ACL force were 58% and 42% in males and 45% and 55% in females at 3 months (Fig. 3B, C). However, AM bundle contribution increased throughout adolescence, and by 18 months, the AM bundle carried 90% of the anterior force in the ACL for both sexes. To statistically compare relative bundle function between sexes and age groups, we calculated the difference in force contribution of the AM bundle minus the force contribution of the PL bundle (Fig. 3D). This difference in bundle contribution increased significantly throughout growth (indicating a shift toward greater AM bundle function) by 19% in males and 40% in females (p<0.001) from 1.5-18 months. Additionally, this metric was greater in males than females by 30%, pooled across all ages (p=0.001). The difference in bundle contribution was statistically significant (p< 0.005 compared to zero, indicating AM bundle contribution > PL bundle contribution) at 1.5, 4.5, 6, and 18 months in males, and at 6 and 18 months in females.

**Figure 3.**
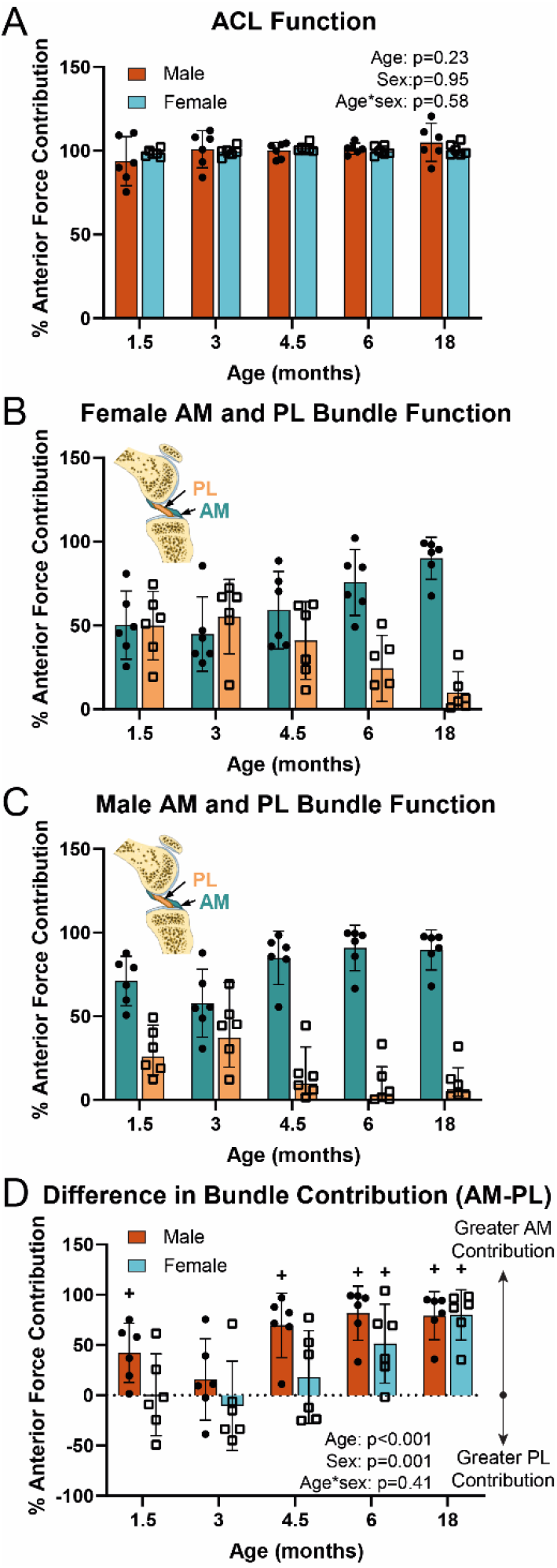
ACL bundle function throughout skeletal growth varied in an age and sex-dependent manner. (A) The ACL contribution to anterior force in the knee joint under maximum anterior translation was nearly 100% in both sexes at all age groups. (B) Female and (C) Male anteromedial (AM) and posterolateral (PL) bundle function diverged throughout growth, such that the AM bundle became dominant by 6 months in females and 4.5 months in males. (D) The difference in bundle contribution (AM bundle % contribution minus PL bundle % contribution) increased throughout skeletal growth and was greater in males than females. Data presented as mean ± 95% confidence interval with main effects from two-way ANOVA shown in graph corner. + indicates p<0.01 relative to zero (indicating AM ≠ PL) within age groups.

### ACL and Bundle Morphology

Mean ACL length increased by approximately 2-fold in males and females from 1.5-18 months (p<0.001, Fig. 4A, Table S9). Mean ACL length was within 1 mm for age matched males and females until 18 months, where the ACL was 15% longer in males than females (p=0.001). Similarly, the AM bundle length increased by 2 and 1.5-fold in males and females, respectively, from 1.5-18 months (p<0.001), and AM bundle length was 11% larger in males than females at 18 months (p=0.01). The PL bundle was consistently shorter than the AM bundle and grew by 2-fold from 1.5-18 months in both sexes (p<0.001). There were no significant differences between sexes within individual age groups, but across all ages, the PL bundle was 5% longer in males than females (p=0.009).

**Figure 4.**
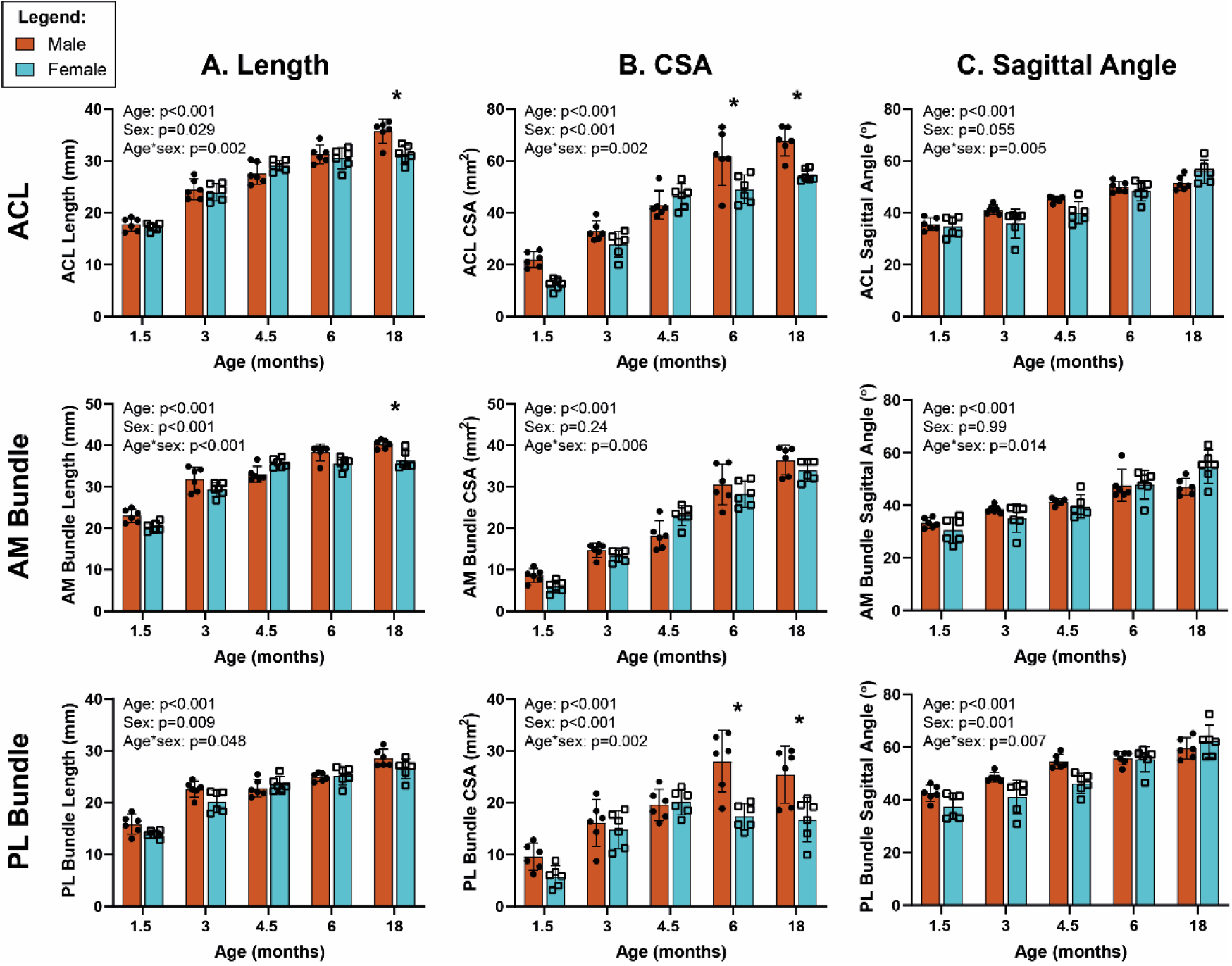
The ACL and its bundles increased in length, CSA, and sagittal angle throughout skeletal growth, in a sex-dependent manner. (A) Length of the ACL and AM bundle were larger in males than females at 18 months, while the PL bundle length remained similar between sexes. (B) Cross-sectional area (CSA) of the ACL and PL bundle were larger in males than females at 6 and 18 months, while the AM bundle CSA remained similar between sexes. (C) Sagittal angle of the ACL, AM bundle, and PL bundle relative to the tibial plateau were similar between males and females throughout skeletal growth. Data presented as mean ± 95% confidence interval with main effects from two-way ANOVA shown in graph corner. * indicates p<0.01 between males and females within age groups.

Mean ACL CSA increased 3 and 4.5-fold in males and females, respectively, from 1.5-18 months (p<0.001, Fig 4B, Table S10). Additionally, the ACL CSA was 23% larger in males than females at both 6 and 18 months (p=0.004 and p=0.002, respectively). AM bundle CSA increased 4 and 5.5-fold in males and females, respectively, from 1.5-18 months (p<0.001). Unlike the full ACL, the AM bundle CSA did not significantly differ between males and females (p=0.24). However, the PL bundle CSA increased 2-3-fold in both sexes (p<0.001) and was 49% (p<0.001) and 38% (p=0.006) larger in males than females at 6 and 18 months, respectively.

Sagittal angle of the ACL relative to the tibial plateau increased by 15° and 21° from 1.5-18 months in males and females, respectively (p<0.001, Fig. 4C, Table S11). Similar increases with age were observed for both the AM and PL bundles (p<0.001). There were not significant differences in sagittal angle of the ACL or AM bundle between males and females. However, the PL bundle was 4° steeper in males than females, across all ages (p=0.001).

### Correlation between Structure and Function

ACL *in situ* stiffness and CSA were positively correlated throughout skeletal growth in males (r^2^=0.75, p<0.001) and females (r^2^=0.64, p<0.001), pooled across all ages (Fig. 5A). However, within age groups, a significant positive correlation was found only for 18 month-old males (r^2^=0.75, p=0.03), with poor fits (0.0003≤r^2^≤0.40) for other age groups (Fig. 5B, Table S13). These seemingly contradictory conclusions demonstrate Simpson’s paradox. AM and PL bundle *in situ* stiffness versus CSA correlations were examined as a function of estimated pubertal status (Fig. 6, Table S14). Correlations for the AM bundle had steeper slopes post-puberty (2.02 in males and 1.82 in females) compared to pre-puberty (1.31 in males and 0.8 in females) and were statistically significant in pre- (r^2^=0.33, p=0.01) and post-pubertal males (r^2^=0.54, p=0.006) and pre-pubertal females (r^2^=0.54, p<0.001). However, correlations for the PL bundle had smaller slopes post-puberty particularly for females (0.89 in males and 0.06 in females) compared to pre-puberty (1.30 in males and 1.24 in females). These correlations were statistically significant only in pre-pubertal males (r^2^=0.59, p<0.001) and females (r^2^=0.54, p<0.001).

**Figure 5.**
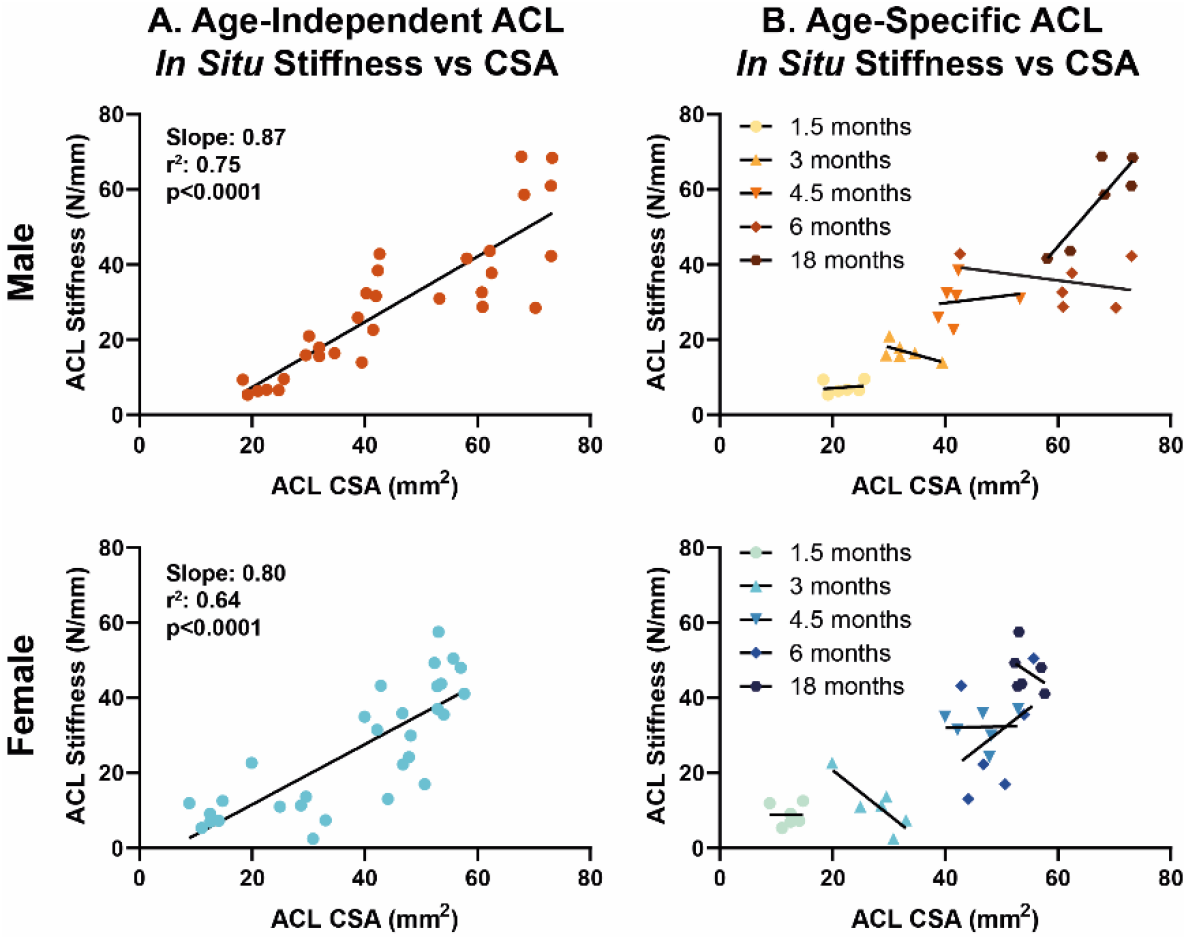
ACL *in situ* stiffness was correlated with ACL cross-sectional area (CSA) throughout skeletal growth but not within individual age groups. (A) Linear regressions for male and female ACL *in situ* stiffness vs CSA, independent of age. Statistical results shown in graph corner. (B) Linear regressions for male and female ACL *in situ* stiffness vs CSA, within age groups. Statistical results reported in Table S13.

**Figure 6.**
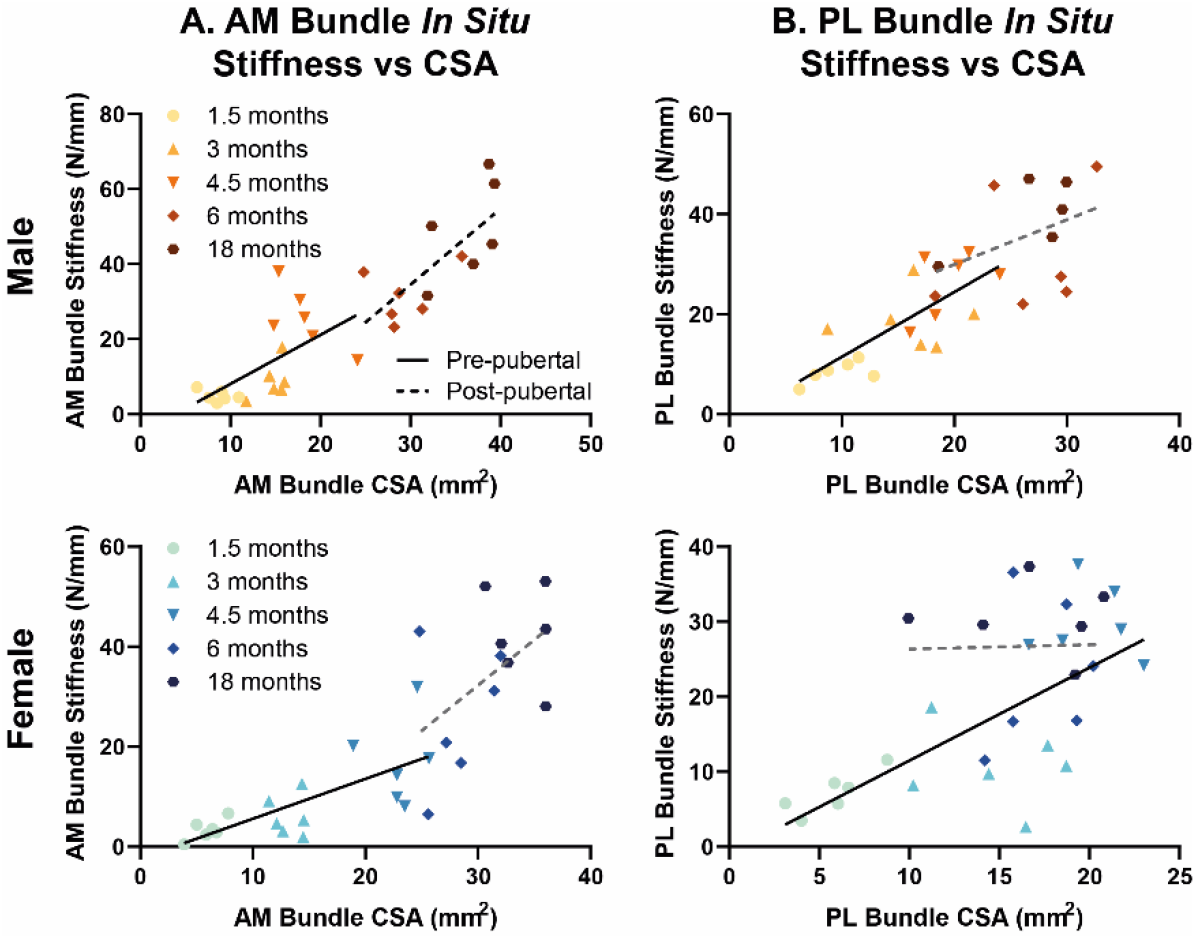
Correlations between *in situ* stiffness and cross-sectional area (CSA) and of the AM and PL bundle are dependent on sex and estimated pubertal status. (A) AM bundle *in situ* stiffness and CSA correlated significantly in pre- and post-pubertal males and pre-pubertal females, with steeper slopes in post-pubertal animals. (B) PL bundle *in situ* stiffness and CSA correlated in pre-pubertal males and females, with smaller slopes in post-pubertal animals, particularly in females. Statistical results of linear regressions reported in Table S14 (black lines indicate p<0.05, gray lines indicate p>0.05).

## Discussion

This study examined the sex-specific function and morphology of the ACL and its bundles throughout skeletal growth in the porcine model. In both males and females, AP joint laxity decreased while joint stiffness increased with increasing age. These changes led to gradual shifts in joint force-translation curves across age groups in females but particularly large shifts from mid to late adolescence in males. The ACL was the primary stabilizer against anterior tibial loading throughout growth in both sexes. But within the ACL, the functional role of the AM bundle increased with age, with this shift occurring earlier in males. The ACL and its bundles increased in length, CSA, and sagittal angle with age. ACL and bundle size were similar in males and females in younger age groups, but ACL and AM bundle length as well as ACL and PL bundle CSA were larger in males than females during adolescence. Additionally, ACL *in situ* stiffness and CSA were correlated throughout skeletal growth, but not necessarily within age groups. AM bundle *in situ* stiffness and CSA were correlated in pre- and post-pubertal males and pre-pubertal females. Finally, PL bundle *in situ* stiffness and CSA were correlated only in pre-pubertal males and females.

The findings with respect to age- and sex-specific biomechanics are generally consistent with human studies.^9–11^ Studies in skeletally immature patients found no difference between sexes during youth, but 20-40% greater translation in females during adolescence.^9; 10^ Similarly, the current study showed that normalized APTT was similar (within 16%) between sexes from 1.5-6 months and 38% greater in females than males at 18 months, though not statistically significant. Some differences may exist since past human studies report raw (not normalized) AP or anterior tibial translation under applied static loads (15-30 lbs) which remained constant across all ages. There are little available human pediatric data regarding function of the ACL and its bundles. However, AM bundle contribution to ACL anterior force in 18 month pigs from this study (90%) is similar to data from adult humans (84%).^31^ Similarly, AM bundle contribution from 6 month pigs in this study (76%-91%) is similar to previously reported data from 6 month pigs (65%).^31^ Additionally, in mature human cadaveric specimens, ACL stiffness under uniaxial tension was 43% lower in females than males,^39^ while in this study, ACL *in situ* stiffness was 19% lower in females at 18 months, though not statistically significant.

The changes in ACL morphology with respect to growth and sex from our study are also consistent with skeletally immature human data. Studies have shown approximately two-fold increases in ACL length and CSA,^6–8^ and approximately 15° increases in angular inclination^8; 33^ from early youth to late adolescence. These studies have also found that while ACL length and CSA are similar between the sexes during childhood, both measurements are 15-25% larger in males during late adolescence.^6–8^ Similarly, our findings showed that ACL length and CSA were 15% and 23% larger, respectively, in males than females during late adolescence. Additionally, in agreement with the current study, human studies found little to no (0-3°) impact of sex on ACL inclination angle.^7; 33^

These data reveal the complex impact of maturation on ACL biomechanics and morphology. Some parameters, such as joint APTT, slack, and stiffness, and ACL size shifted considerably in males from 6 to 18 months (mid to late adolescence), with smaller or no changes in females. This may indicate earlier maturation in females. However, looking at ACL bundle function, there was an earlier shift to AM bundle dominance in males (3-4.5 months) than females (4.5-6 months). This would seem to indicate later maturation in females. Adding to the complexity, we observed sex-specific changes in ACL bundle size during adolescence. Specifically, male and female AM bundles were comparable in CSA at all ages while the PL bundle grew larger in males than females during adolescence, despite carrying less force under loading. This was surprising, since we expected differences in bundle function between sexes to correspond to differences in bundle size. This seems to indicate that other biological factors, such as hormones, could contribute to sex-specific ACL growth, and may play a role in decoupling changes in size from changes in function in some cases, which is evident in the lack of correlation between *in situ* stiffness and CSA for the post-pubertal PL bundle, particularly for female animals.

Results from this study are relevant to understanding age and sex-dependent ACL injury risk since smaller ACL size has been identified as an injury risk factor,^13^ and we found that the female ACL is smaller than the male ACL during adolescence (6-18 months). Additionally, joint laxity has been identified as an injury risk factor in adults,^12–14^ and our results showed that AP stifle joint laxity trended higher in females than males during late adolescence. However, this difference in joint laxity is small in magnitude and depends on several underlying factors including ACL and bundle size, orientation, and mechanical properties. In fact, a recent study in humans found that sex and weight did not impact anterior knee laxity after accounting for ACL volume and T2* relaxation time.^40^ Therefore, evaluating the effect of underlying factors, such as bundle-specific properties and *in situ* slack and stiffness, on ACL injury risk may help us better understand age and sex-dependent injury rates. Although not measured in this study, there is also evidence that bony anatomical features,^13; 14^ neuromuscular biomechanics,^41–43^ sex hormones,^44–46^ and oral contraceptive use^46–48^ impact ACL injury risk and likely age and sex-dependent injury rates. Understanding the interaction between these many risk factors may be key to preventing ACL injuries in young patients.

Our findings are also relevant to ACL injury treatment in the pediatric population. For example, given rapid changes in ACL size, orientation, and bundle function during skeletal growth, how should graft size and tunnel locations be selected in pediatric patients? Our findings seem to support the usefulness of approaches that can appropriately grow and mature with the patient. It is possible that biologically enhanced suture repair techniques^26; 49^ may be more effective in this regard. Additionally, this work shows that ACL *in situ* stiffness can be estimated as a function of CSA throughout skeletal growth. Given the paucity of data regarding the biomechanical properties of the human pediatric ACL, ligament area measured from clinical MRI scans may be useful to estimate changes in biomechanical properties across age groups. However, it is important to note that these correlations only held true over the full scale of skeletal growth, and not within specimens of the same age. Overall, our findings indicate that the pediatric ACL is not simply a scaled down version of the skeletally mature ACL, due to age- and sex-specific differences in the size and function of the AM and PL bundles, and accounting for these differences may benefit clinical outcomes.

There are several limitations of this study. Firstly, although the pig is a good model for the human ACL, some anatomical differences exist. Primarily, the AM bundle inserts anterior to the lateral meniscal root on the tibia in pigs but not in humans. This difference may cause subtle differences in geometry and function. Additionally, porcine stifle joints have a smaller range of joint flexion than human knees (porcine stifle full extension at ~40°), and therefore native loading conditions of the ACL and its bundles may differ slightly between pigs and humans. Additionally, it was not feasible to measure animal weight, which is an important clinical factor. Furthermore, the biomechanical tests performed in this study simulate clinical anterior drawer exams rather than *in vivo* motions. Therefore, it is likely that this work does not completely capture the *in vivo* loading patterns of the ACL. Finally, this is a cross-sectional study and does not capture growth within individual subjects.

In conclusion, this study found age and sex-specific morphological and functional maturation of the ACL throughout skeletal growth. Specifically, the functional role of the AM bundle increased throughout growth in both sexes but became dominant over the PL bundle earlier in males compared to females. During adolescence, the ACL and PL bundle CSA and the ACL and AM bundle length were larger in males than females. Finally, we found that for the ACL, *in situ* stiffness and CSA correlated throughout skeletal growth, while for the AM and PL bundles, correlations differed as a function of pubertal status. This study highlights the potential need to account for age- and sex-specific changes within the ACL to improve treatment of pediatric ACL injuries.

## Acknowledgments

We would like to thank Joella Knopf, BS, Lauryn Braxton, BS, and Stephanie Teeter, MA, for their assistance during robotic testing experiments and Laura Edwards, BS, for her assistance with animal work. Additionally, we would like to thank the Swine Education Unit at North Carolina State University and the Biomedical Research Imaging Center at the University of North Carolina- Chapel Hill for their contributions to this work. Research in this publication was supported by the National Institute of Arthritis and Musculoskeletal and Skin Diseases of the U.S. National Institutes of Health (R01AR071985, R03AR068112). The content is solely the responsibility of the authors and does not necessarily represent the official views of the National Institutes of Health. Additional funding for this study was provided in part by the National Science Foundation (DGE-1252376).

## Notes

### Competing Interest Statement

The authors have declared no competing interest.

