## Supplemental Material for "Sex-Specific Function and Morphology of the Anterior Cruciate Ligament During Skeletal Growth in a Porcine Model"

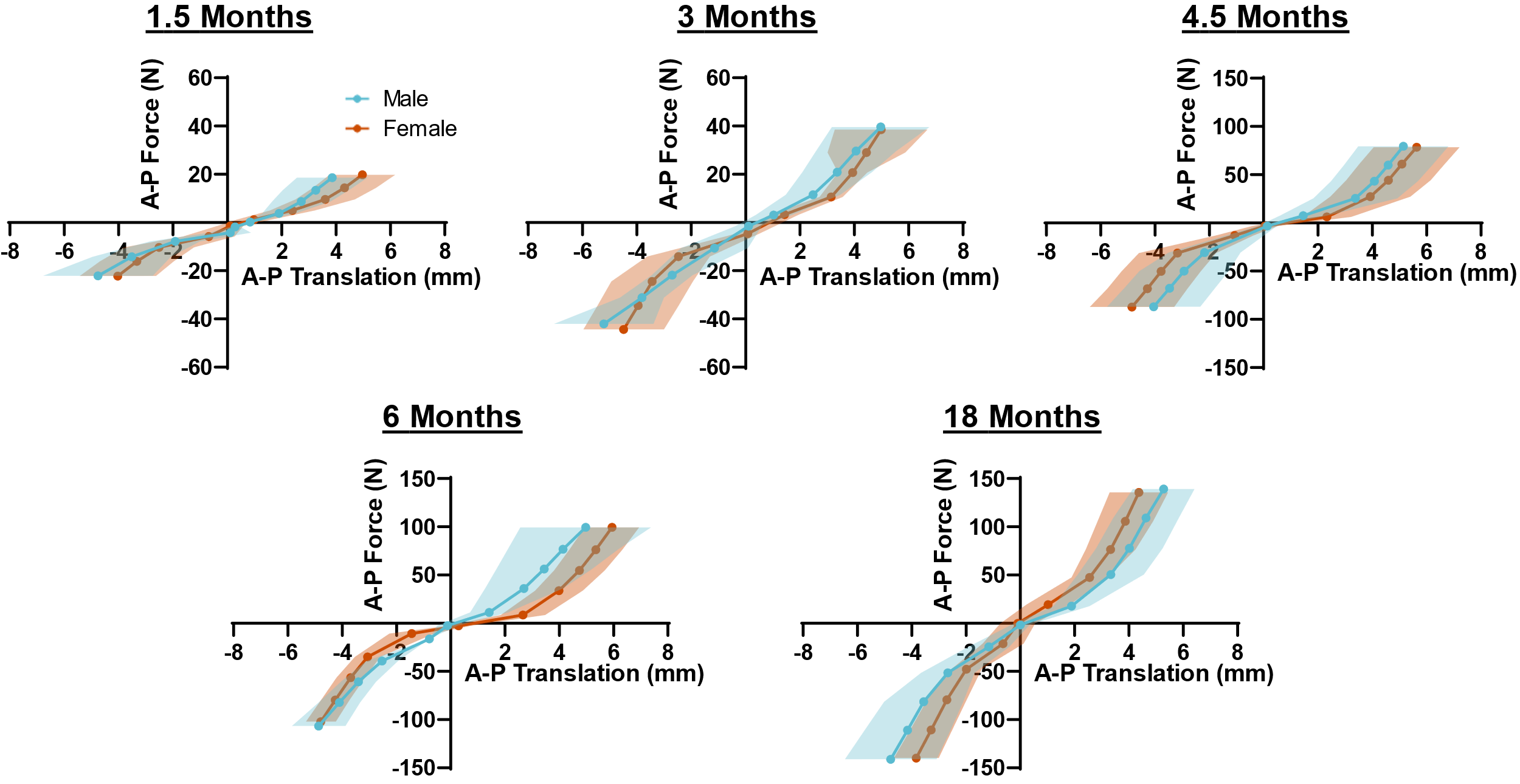

**Figure S1.** Mean force-translation curves comparing male and female joints within each age group (shadow indicates standard deviation in translation, interpolated between each data point). The force-translation curves appeared similar between sexes at 1.5, 3, and 4.5 months. The average male force-translation curve extended out wider than females at 6 months, but was steeper and narrower at 18 months.

**Table S1.** Preliminary APTT data from female joints with and without locked internal rotation.

| APTT (mm) | | |
| --- | --- | --- |
| Age | IR Locked | IR Unlocked |
| 3 months | 7.79 ± 0.87 | 7.82 ± 0.83 |
| 4.5 months | 10.21 ± 1.53 | 10.21 ± 1.54 |
| 6 months | 8.18 ± 1.72 | 8.10 ± 1.77 |
| 18 months | 8.58 ± 0.73 | 8.58 ± 0.75 |

Data is presented as mean ± SD. IR: internal rotation.

**Table S2.** Preliminary force contribution data of the capsule under anterior drawer.

| Capsule Contribution | |
| --- | --- |
| Age | Force (N) |
| 3 months | 2.1 ± 1.5 |
| 4.5 months | 0.3 ± 2.5 |
| 6 months | 1.4 ± 1.9 |
| 18 months | 0.1 ± 2.8 |

Data is presented as mean ± SD.

**Table S3**. Normalized APTT Summary Data

| Normalized APTT | | |
| --- | --- | --- |
| Age | Male | Female |
| 1.5 months | 0.26 ± 0.01 | 0.28 ± 0.06 |
| 3 months | 0.24 ± 0.02 | 0.24 ± 0.07 |
| 4.5 months | 0.21 ± 0.03 | 0.19 ± 0.03 |
| 6 months | 0.20 ± 0.02 | 0.17 ± 0.05 |
| 18 months | 0.13 ± 0.01 | 0.19 ± 0.02 |

Data is presented as mean ± SD. APTT: anterior-posterior tibial translation.

**Table S4**. Raw APTT Summary Data

| APTT (mm) | | |
| --- | --- | --- |
| Age | Male | Female |
| 1.5 months | 9.0 ± 0.7 | 8.6 ± 1.5 |
| 3 months | 9.5 ± 0.7 | 10.3 ± 3.4 |
| 4.5 months | 10.5 ± 1.1 | 9.2 ± 1.3 |
| 6 months | 10.7 ± 0.9 | 9.8 ± 3.0 |
| 18 months | 8.2 ± 1.2 | 10.1 ± 0.9 |

Data is presented as mean ± SD. APTT: anterior-posterior tibial translation.

**Table S5**. Joint *In Situ* Slack Length Summary Data

| Joint *In Situ* Slack (mm) | | |
| --- | --- | --- |
| Age | Male | Female |
| 1.5 months | 5.2 ± 0.7 | 3.8 ± 0.7 |
| 3 months | 5.5 ± 0.8 | 3.8 ± 1.4 |
| 4.5 months | 6.2 ± 1.0 | 5.2 ± 1.1 |
| 6 months | 5.9 ± 0.8 | 4.5 ± 1.0 |
| 18 months | 4.2 ± 0.8 | 5.4 ± 0.8 |

Data is presented as mean ± SD.

**Table S6**. Joint *In Situ* Stiffness Summary Data

| *In Situ* Joint Stiffness (N/mm) | | |
| --- | --- | --- |
| Age | Male | Female |
| 1.5 months | 7.2 ± 1.4 | 9.2 ± 2.7 |
| 3 months | 15.6 ± 3.7 | 13.4 ± 5.1 |
| 4.5 months | 29.0 ± 5.9 | 31.3 ± 5.7 |
| 6 months | 33.8 ± 6.1 | 34.8 ± 14.7 |
| 18 months | 52.0 ± 13.2 | 44.9 ± 4.1 |

Data is presented as mean ± SD.

**Table S7**. ACL Contribution to Anterior Drawer

| ACL Contribution (% of Joint Anterior Force) | | |
| --- | --- | --- |
| Age | Male | Female |
| 1.5 months | 94 ± 14 | 99 ± 2 |
| 3 months | 101 ± 11 | 99 ± 3 |
| 4.5 months | 100 ± 5 | 102 ± 2 |
| 6 months | 101 ± 3 | 100 ± 3 |
| 18 months | 105 ± 11 | 101 ± 4 |

Data is presented as mean ± SD.

**Table S8**. AM and PL Bundle Contribution to ACL Anterior Drawer

|  | AM Bundle Contribution  (% of ACL Anterior Force) | | PL Bundle Contribution  (% of ACL Anterior Force) | |
| --- | --- | --- | --- | --- |
| Age | Male | Female | Male | Female |
| 1.5 months | 71 ± 14 | 50 ± 19 | 29 ± 14 | 50 ± 19 |
| 3 months | 58 ± 19 | 45 ± 21 | 42± 19 | 55 ± 21 |
| 4.5 months | 85 ± 15 | 59 ± 22 | 15 ± 15 | 41 ± 22 |
| 6 months | 91 ± 13 | 76 ± 19 | 9 ± 13 | 24 ± 19 |
| 18 months | 90 ± 11 | 90 ± 12 | 10 ± 11 | 10 ± 12 |

Data is presented as mean ± SD.

**Table S9**. ACL, AM and PL Bundle Length

|  | ACL Length (mm) | | AM Bundle Length (mm) | | PL Bundle Length (mm) | |
| --- | --- | --- | --- | --- | --- | --- |
| Age | Male | Female | Male | Female | Male | Female |
| 1.5 months | 17 ± 1 | 17 ± 1 | 23 ± 2 | 20 ± 1 | 16 ± 2 | 14 ± 1 |
| 3 months | 25 ± 2 | 24 ± 2 | 32 ± 3 | 29 ± 2 | 23 ± 1 | 20 ± 2 |
| 4.5 months | 28 ± 2 | 29 ± 1 | 33 ± 2 | 36 ± 1 | 23 ± 2 | 23 ± 1 |
| 6 months | 31 ± 2 | 31 ± 2 | 38 ± 2 | 36 ± 1 | 25 ± 1 | 25 ± 2 |
| 18 months | 36 ± 2 | 31 ± 2 | 40 ± 1 | 36 ± 2 | 29 ± 2 | 27 ± 2 |

Data is presented as mean ± SD.

**Table S10**. ACL, AM and PL Bundle Cross-Sectional Area (CSA)

|  | ACL CSA (mm^2^) | | AM Bundle CSA (mm^2^) | | PL Bundle CSA (mm^2^) | |
| --- | --- | --- | --- | --- | --- | --- |
| Age | Male | Female | Male | Female | Male | Female |
| 1.5 months | 22 ± 3 | 12 ± 2 | 9 ± 2 | 6 ± 1 | 10 ± 2 | 6 ± 2 |
| 3 months | 32 ± 4 | 28 ± 5 | 15 ± 2 | 13 ± 1 | 16 ± 4 | 15 ± 3 |
| 4.5 months | 43 ± 5 | 46 ± 5 | 18 ± 3 | 23 ± 2 | 20 ± 3 | 20 ± 2 |
| 6 months | 62 ± 11 | 49 ± 5 | 31 ± 5 | 28 ± 3 | 28 ± 6 | 17 ± 2 |
| 18 months | 68 ± 6 | 54 ± 2 | 36 ± 3 | 34 ± 2 | 25 ± 5 | 17 ± 4 |

Data is presented as mean ± SD.

**Table S11**. ACL, AM and PL Bundle Sagittal Angle

|  | ACL Sagittal Angle (°) | | AM Bundle Sagittal Angle (°) | | PL Bundle Sagittal Angle (°) | |
| --- | --- | --- | --- | --- | --- | --- |
| Age | Male | Female | Male | Female | Male | Female |
| 1.5 months | 36 ± 2 | 35 ± 4 | 33 ± 2 | 30 ± 5 | 42 ± 3 | 37 ± 5 |
| 3 months | 41 ± 2 | 36 ± 5 | 38 ± 1 | 35 ± 5 | 48 ± 2 | 41 ± 6 |
| 4.5 months | 45 ± 1 | 40 ± 4 | 41 ± 1 | 40 ± 4 | 55 ± 3 | 46 ± 4 |
| 6 months | 50 ± 2 | 48 ± 4 | 48 ± 6 | 48 ± 5 | 56 ± 2 | 55 ± 5 |
| 18 months | 51 ± 3 | 56 ± 4 | 47 ± 3 | 55 ± 6 | 59 ± 4 | 62 ± 6 |

Data is presented as mean ± SD.

**Table S12**. ACL, AM and PL Bundle *In Situ* Stiffness

|  | ACL *In Situ* Stiffness (N/mm) | | AM Bundle *In Situ* Stiffness (N/mm) | | PL Bundle *In Situ* Stiffness (N/mm) | |
| --- | --- | --- | --- | --- | --- | --- |
| Age | Male | Female | Male | Female | Male | Female |
| 1.5 months | 7 ± 2 | 9 ± 3 | 5 ± 1 | 3 ± 2 | 8 ± 2 | 7 ± 3 |
| 3 months | 17 ± 2 | 11 ± 7 | 9 ± 5 | 6 ± 4 | 19 ± 6 | 11 ± 5 |
| 4.5 months | 30 ± 5 | 32 ± 5 | 26 ± 8 | 17 ± 9 | 26 ± 7 | 30 ± 5 |
| 6 months | 35 ± 6 | 30 ± 15 | 32 ± 7 | 26 ± 14 | 32 ± 12 | 23 ± 10 |
| 18 months | 57 ± 12 | 47 ± 6 | 49 ± 13 | 42 ± 9 | 40 ± 7 | 31 ± 5 |

Data is presented as mean ± SD.

**Table S13**. ACL *In Situ* Stiffness vs Cross-Sectional Area—Age-Specific Linear Regression Results

|  | Male | | | Female | | |
| --- | --- | --- | --- | --- | --- | --- |
| Age | Slope | r^2^ | p-value | Slope | r^2^ | p-value |
| 1.5 months | 0.11 | 0.03 | 0.73 | -0.02 | 0.0003 | 0.97 |
| 3 months | -0.42 | 0.40 | 0.18 | -1.19 | 0.69 | 0.04 |
| 4.5 months | 0.18 | 0.03 | 0.74 | 0.03 | 0.001 | 0.96 |
| 6 months | -0.20 | 0.11 | 0.53 | 1.17 | 0.17 | 0.42 |
| 18 months | 1.71 | 0.75 | 0.03 | -1.00 | 0.15 | 0.45 |

**Table S14.** AM and PL Bundle *In Situ* Stiffness vs Cross-Sectional Area—Pre- and Post-Pubertal Linear Regression Results

|  | AM Bundle | | | PL Bundle | | |
| --- | --- | --- | --- | --- | --- | --- |
| Age | Slope | r^2^ | p-value | Slope | r^2^ | p-value |
| Male: Pre-pubertal | 1.31 | 0.33 | 0.01 | 1.30 | 0.59 | <0.001 |
| Male: Post-pubertal | 2.02 | 0.54 | 0.006 | 0.89 | 0.16 | 0.23 |
| Female: Pre-pubertal | 0.80 | 0.54 | <0.001 | 1.24 | 0.54 | <0.001 |
| Female: Post-pubertal | 1.82 | 0.25 | 0.09 | 0.06 | 0.001 | 0.94 |
